# Haplotypes spanning centromeric regions reveal persistence of large blocks of archaic DNA

**DOI:** 10.1101/351569

**Authors:** Sasha A. Langley, Karen Miga, Gary Karpen, Charles H. Langley

## Abstract

Despite critical roles in chromosome segregation and disease, the repetitive structure and vast size of centromeres and their surrounding heterochromatic regions impede studies of genomic variation. We report here large-scale haplotypes (*cenhaps*) in humans that span the centromere-proximal regions of all metacentric chromosomes, including the arrays of highly repeated α-satellites on which centromeres form. *Cenhaps* reveal surprisingly deep diversity, including entire introgressed Neanderthal centromeres and equally ancient lineages among Africans. These centromere-spanning haplotypes contain variants, including large differences in α-satellite DNA content, which may influence the fidelity and bias of chromosome transmission. The discovery of *cenhaps* creates new opportunities to investigate their contribution to phenotypic variation, especially in meiosis and mitosis, as well as to more incisively model the unexpectedly rich evolution of these challenging genomic regions.

**One Sentence Summary:** Genomic polymorphism across centromeric regions of humans is organized into large-scale haplotypes with great diversity, including entire Neanderthal centromeres.

---

The centromere is the unique chromosomal locus that forms the kinetochore, which interacts with spindle microtubules and directs segregation of replicated chromosomes to daughter cells (*1*). Human centromeres assemble on a subset of large blocks (many Mbps) of highly repeated (171 bp) α-satellite arrays found on all chromosomes. These repetitive arrays and the flanking segments, together the Centromere Proximal Regions (CPRs), play critical roles in the integrity of mitotic and meiotic inheritance (*2*). In somatic tissues, chromosome instability, including loss and gain of chromosomes, plays large and complex roles in aging, cancer (*3*), and human embryonic survival (*4*). Sequence variation in CPRs can affect meiotic pairing (*3,4*), kinetochore formation (*5,6*) and nonrandom segregation (*3,4,6*). Aneuploidy in the germline, typically arising during meiosis, is a large component of genetic disease (*9*). Further, the unique asymmetry of transmission in female meiosis, where only one parental chromosome is transmitted, presents the opportunity for the evolution of strong deviations from mendelian segregation ratios (meiotic drive) (*10,11*). The large selective impact of recurrent meiotic drive is one potential cause of the evolutionarily rapid divergence of satellite DNAs and centromeric chromatin proteins (*12*), reduced polymorphisms in flanking regions and high levels of aneuploidy (*13*). The challenges inherent to assessing genomic variation in these repetitive and dynamic regions remain a significant barrier to incisive functional and evolutionary investigations.

Recognizing the potential research value of well-genotyped diversity across human CPRs, we hypothesized that the low rates of meiotic exchange in these regions (*14*) might result in large, haplotypes in populations, perhaps even spanning both the α-satellite arrays. To test this, we examined the Single Nucleotide Polymorphism (SNP) linkage disequilibrium (LD) and haplotype variation surrounding the centromeres among the diverse collection of genotyped individuals in Phase 3 of the 1000 Genomes Project (*15*). Figure 1a depicts the predicted patterns of strong LD (red) and associated unbroken haplotypic structures surrounding the gap of unassembled satellite DNA of a metacentric chromosome. Unweighted Pair Group Method with Arithmetic Mean (UMPGA) clustering on 800 SNPs immediately flanking the chrX centromeric gap in males (Fig. 1c) reveals a clear haplotypic structure that spans the gap and extends, as predicted, to a much larger region (≈7 Mbp, Fig. 1b). Similar clustering of the imputed genotypes of females also falls into the same distinct high-level haplotypes (Fig. S1). This discovery of the predicted haplotypes spanning CPRs (hereafter referred to as *cenhaps*) on chrX and all the metacentric chromosomes (Fig. S2) opens a new window into their evolutionary history and functional potential.

**Figure 1.**
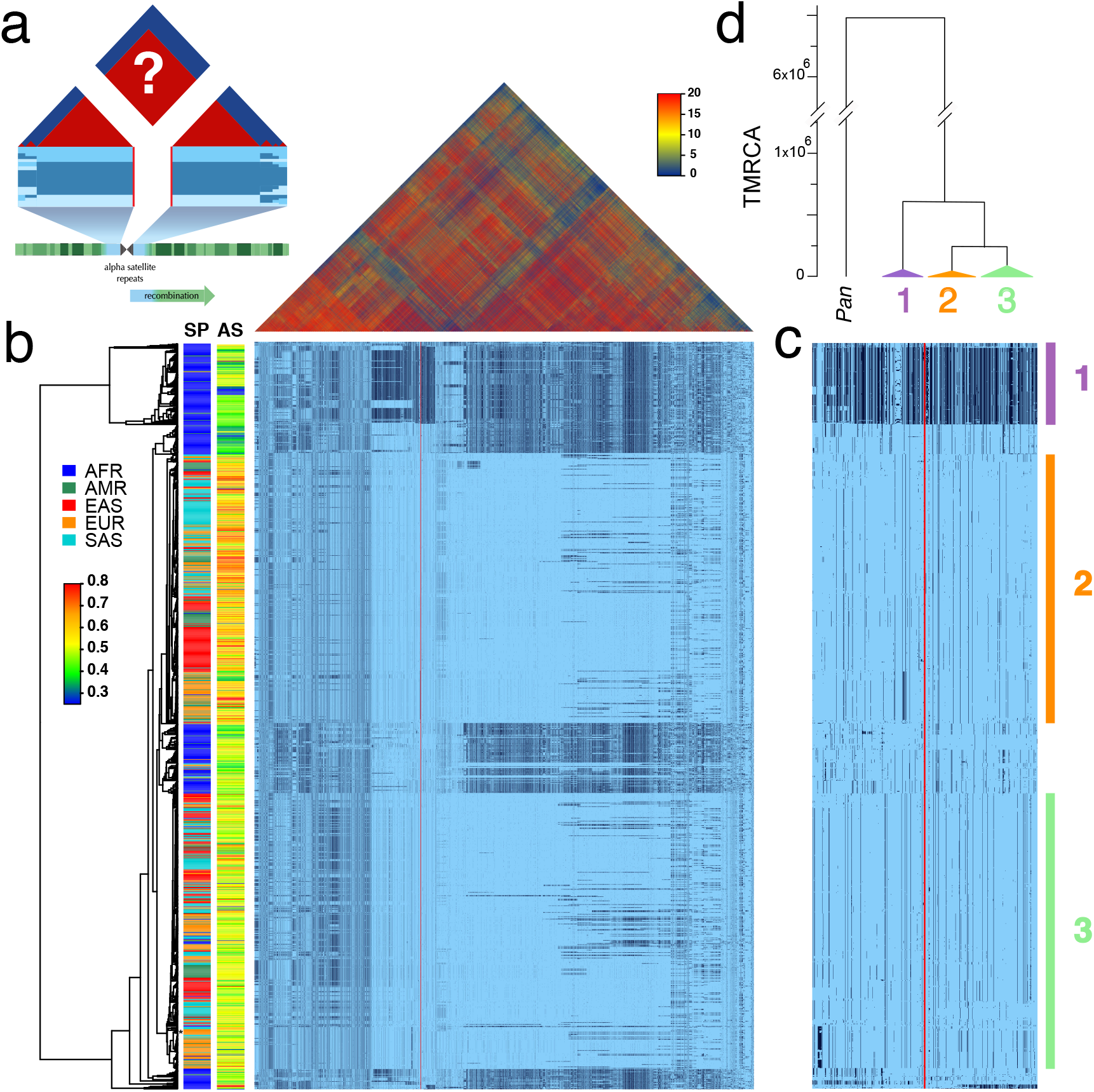
Strong LD across centromeric gaps forms large-scale centromere-spanning haplotypes, or *cenhaps.* **a**. The predicted patterns of the magnitude LD (triangle at top) and genotypes in CPR clustered into haplotypes surrounding the centromeric region of a metacentric chromosome in a large outbreeding population (central blue bands) if crossing over declines to zero in and around the highly repeated DNA where the centromere is typically found in the chromosome (blue and green at bottom of **a**). **b**. Above, the linkage disequilibria between pairs of 17702 SNPs (Left: chrX:55623011-58563685, Right: chrX: 61725513-68381787; hg19) flanking the centromere and α-satellite assembly gap (red vertical line) from 1231 human male X chromosomes from the 1000 Genomes Project. The color maps (see the adjacent legend) to the -*log_10_(p)* where the *p* value derives from the 2×2 ***χ***^2^ for each pair of SNPs. Below, a broad haplotypic representation of these same data. SNPs were filtered for minor allele count (MAC)≥60. Minor alleles shown in black. Poorly genotyped SNPs near edges of the gap (red line) were masked. Superpopulation (**SP**; **AFR**ica, **AMeR**icas, **E**ast **AS**ia, **EUR**ope, **S**outh **AS**ia) and scaled estimate of chrX-specific α-satellite array size (**AS**) indicated at left side. Approximate position of HuRef chrX indicated by black asterisk at right of the tree. Dendrogram represents UPGMA clustering based on the hamming distance between haplotypes comprised of 800 filtered SNPs immediately flanking the centromere (Left: chrX:58374895-58563685, Right: chrX:61725513-61921419; hg19), shown in detail in c. The three most common X cenhaps highlighted with colored vertical bars. **d**. A UPGMA tree based on the synonymous divergence in 17 genes (see Table S1) in the 3 major chrX cenhaps (indicated in c), assuming the TMRCA of humans and chimps is 6.5MY. Widths of the triangles are proportional to the *log_10_* of number of members of each cenhap, and the height is proportional to the average divergence within each cenhap.

The pattern of geographic differentiation across the inferred chrX CPR (Fig. 1) exhibits higher diversity in African samples, as observed throughout the genome (*15*). Despite being fairly common among Africans today, a distinctly diverged chrX cenhap (cenhap 1, highlighted in purple, Fig. 1b,c) is rare outside of Africa. Examination of the haplotypic clustering and estimated synonymous divergence in the coding regions of 21 genes included in the chrX cenhap region (see Table S1) yields a parallel relationship among the three major cenhaps and an estimated Time of the Most Recent Common Ancestor (TMRCA) of ≈700 KYA (Fig. 1d) for this most diverged example. While ancient, putatively introgressed archaic segments have been inferred in African genomes (*16,17*), this cenhap stands out as genomically (if not genetically) large. The persistence of such ancient cenhaps is inconsistent with the predicted hitchhiking effect of sequential fixation of new meiotically driven centromeres (*12*). Further, the detection of near-ancient segments spanning the centromere contrasts with the observation of substantially more recent ancestry across the remainder of chrX and with the expectation of reduced archaic sequences on chrX (*18*). A large block on the right in Fig. 1b, where recombination has substantially degraded the haplotypic structure, is comprised of SNPs in exceptionally high frequency in Africans. Its history in “anatomically modern humans” (AMH) may be shared with the ancient cenhap in Africa. Many distal recombinants are observed outside of Africa that likely contribute to associations of SNPs in this region with a diverse set of phenotypes, including male pattern hair loss and prostate cancer (*19,20*).

This deep history of the chrX CPR raises the possibility of even more ancient lineages on other chromosomes, either derived by admixture with archaic hominins or maintained by balancing forces. A survey of the other chromosomes uncovered several interesting examples (see Fig. S2), two of which we present in detail. To identify Neanderthal and Denisovan introgressed segments we looked for highly diverged alleles in non-African populations that shared a strong excess of derived alleles with archaic hominids, and not with AMH genomes (using chimpanzee as the outgroup; *21,22,23).* Applying this approach to CPR of chr11 revealed a compelling example of Neanderthal introgression, which is illustrated in Fig. 2a in the context of the seven most common chr11 cenhaps. The most diverged lineage contains a small basal group of out-of-Africa genomes (cenhap 1, highlighted in green). Members of this cenhap carry a large proportion of the derived alleles assigned to the Neanderthal lineage, DM/(DM+DN) = 0.98, where DM is the cenhap mean number of shared Neanderthal Derived Matches and DN is the cenhap mean number of Neanderthal Derived Non-matches (Fig2a, at left). The ratio DM/(DM+AN) = 0.91, where AN is the number of Neanderthal-cenhap Non-matches that are Ancestral in the Neanderthal, is a measure of the proportion of the cenhap lineage shared with Neanderthals and further supports the conclusion that this chr11 cenhap is an introgressed archaic centromere. Fig. 2b shows these mean counts for each SNP class by cenhap group, confirming that the affinity to Neanderthals is slightly stronger than to Denisovans. A second basal African lineage separates shortly after the Neanderthal (cenhap 2, highlighted in purple, Fig. 2a). It is unclear if this cenhap represents an introgression from a distinct archaic hominin in Africa or a surviving ancient lineage within the population that gave rise to AMHs.

**Figure 2.**
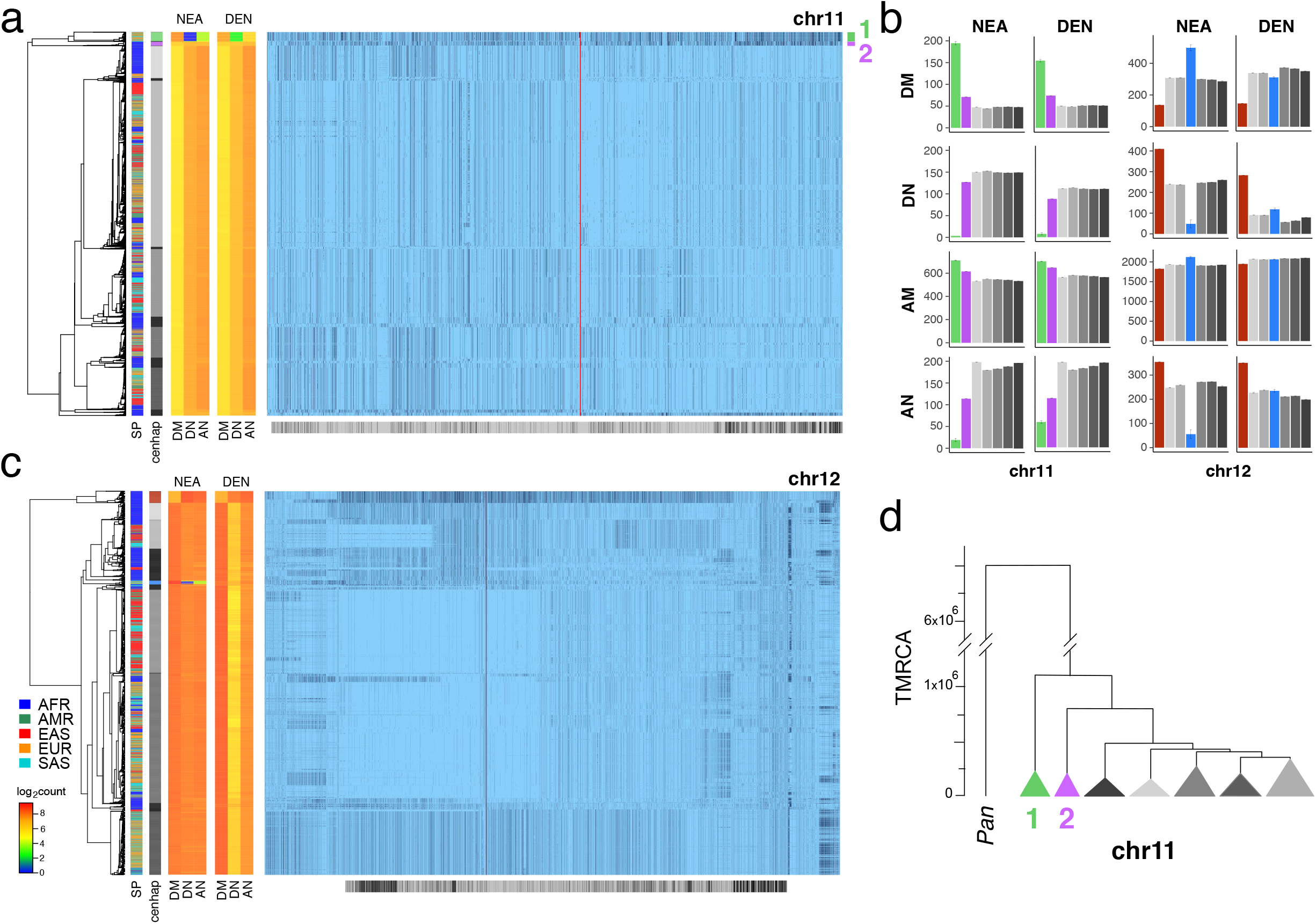
Archaic cenhaps are found in AMH populations. **a**. Haplotypic representation of 9151 SNPs from 5008 imputed chr11 genotypes from the 1000 Genomes Project (Left: chr11:50509493-51594084, Right: chr11:54697078-55326684; hg19). SNPs were filtered for MAC≥35 and passing the *4gt_dco* with a tolerance of three (see Methods). Minor alleles shown in black and assembly gap indicated by red line. Haplotypes were clustered with UPGMA based on the hamming distance between haplotypes comprised of 1000 SNPs surrounding the gap (Left: chr11: 51532172-51594084, Right: chr11:54697078-54845667; hg19). Superpopulation and cenhap partitioning indicated in bars at far left. Log2 counts of DM (derived in archaic, shared by haplotype), DN (derived in archaic, not shared by haplotype) and AN (ancestral in archaic, not shared by haplotype) for each cenhap relative to Altai Neanderthal (NEA) and Denisovan (DEN) at left. Grey horizontal bar (bottom) indicates region included in analysis of archaic content; black bars indicate SNPs with data for archaic and ancestral states. **b**. Bar plots indicating the mean and 95% confidence intervals of DM, DN, AM (ancestral in archaic, shared by cenhap) and AN counts for cenhap groups (as partitioned in a. and c.) relative to Altai Neanderthal and Denisovan genomes, using chimpanzee as an outgroup (23). **c**. Haplotypic representation, as above, of 21950 SNPs from 5008 imputed chr12 genotypes from the 1000 Genomes Project (Left: chr12:33939700-34856380, Right: chr12:37856765-39471374; hg19). SNPs were filtered for MAC≥35. Haplotypes were clustered with UPGMA based on 1000 SNPs surrounding the gap (Left: chr12: 34821738-34856670, Right: chr12:37856765-37923684; hg19). Bars at side and bottom same as in **a. d**. A UPGMA tree based on the synonymous divergence in 30 genes in the 7 major chr11 cenhaps (see Table S2), assuming the TMRCA of humans and chimpanzee is 6.5MY (see Methods and legend for Fig 1d).

The relatively large expanses of these cenhaps and unexpectedly sparse evidence of recombination could be explained by either relatively recent introgressions or cenhap-specific suppression of crossing over with other AMH genomes in this CPR (e.g., an inversion). As with chrX above, the clustering of cenhaps based on coding synonymous SNPs (Fig. 2d) yields a congruent topology and estimates of TMRCAs for the two basal cenhaps of 1.1 and 0.8 MYA, consistent with relatively ancient origins. Among the 37 genes ‘captured’ in this Neanderthal cenhap (1) are 34 of the ~300 known (*24*) odorant receptors (ORs). SNPs in these chr11 ORs are associated with variation in human olfactory perception of particular volatile chemicals (*25*). Given the large number of substitutions, 52 amino acid replacements among 20 of these ORs (Table S2), this cenhap likely codes for Neanderthal-specific determinants of smell and taste. Similarly, in the second ancient cenhap found primarily within Africa (2), eight of these ORs harbor 14 amino acid replacements, of which only two are shared with cenhap 1 (see Table S2). The frequencies of the Neanderthal cenhap in Europe, South Asia and the Americas (0.061, 0.032 and 0.033, respectively), and of second ancient cenhap in Africa (0.036), are sufficiently high that together they contribute more than half of the amino acid replacement diversity in these 34 ORs among the 1000 Genomes (see Table S5). Thus, a substantial part of the variation in chemical perception among AMH may be contributed by these two ancient cenhaps.

The most diverged, basal clade on chr12 (Fig. 2c, indicated in brown) is common in Africa, but, like the most diverged chrX cenhap, is not represented among the descendants of the out-of-Africa migrations (*26*). The great depth of the lineage of this cenhap is further supported by comparison to homologous archaic sequences (*21,22,23*). Consistent with the hypothesis that this branch split off before that of Neanderthals/Denisovans, members of this cenhap share fewer matches with derived SNPs on the Neanderthal and Denisovan lineages (DM) and exhibit strikingly more ancestral non-matches (AN) than other chr12 cenhaps (see Fig. 2b). This putatively archaic chr12 cenhap represents a large and obvious example of the genome-wide introgressions into African populations inferred from model-based analyses of the distributions of sequence divergence (*16,17*). The small out-of-Africa cenhap nested within a mostly African subclade (indicated in blue in Fig. 2c) appears to be a typical Eurasian archaic introgression with higher affinity to Neanderthals (DM/(DN+DM) = 0.91 and DM/(DM+AN) = 0.90) than to Denisovans (Fig. 2b). This bolsters the conclusion that the basal African cenhap represents a distinct, older and likely introgressed archaic lineage. Unfortunately, there are too few coding bases in this region to support confident estimation of the TMRCAs of these ancient chr12 cenhaps. Based on the numbers of SNPs underlying the cenhaps, this basal cenhap is twice as diverged as the apparent introgressed Neanderthal cenhap, placing the TMRCA at ~1.1 MYA, assuming the Neanderthal TMRCA was 575KYA(*23*). While there is no direct evidence of recent introgression, the large genomic scale of this most diverged cenhap (relative to apparent exchanges in other cenhaps) is consistent with recent admixture with an extinct archaic in Africa, although, again, suppression of crossing over is an alternative explanation.

Chromosomes X, 11 and 12 harbor a diversity of large cenhaps, including those representing archaic lineages. Notably, the CPRs of other chromosomes include diverged, basal lineages that are likely to be relatively old, if not archaic (Fig. S2). For example, chromosome 8 contains a ancient cenhap limited to Africa with an estimated TMRCA of 817 KYA (Fig. S3) and a basal chr10 cenhap appears to be another clear Neanderthal introgression (Fig. S4). Obtaining genomic sequence from African archaics would shed light on the evolutionary origins of the ancient cenhaps not associated with Neanderthal and Denisovan introgression. It should be noted that the very large genomic sizes of these ancient cenhaps could allow identification of archaic homology even with modest genomic sequence coverages from archaic fossils.

These SNP-based cenhaps portray a rich, highly structured view of the diversity in the unique segments flanking repetitive regions. While the divergence of satellites may be dynamic on a shorter time scale(*27*), we reasoned that the paucity of exchange in these regions would create cenhap associations with satellite divergence in both sequence and array size. Miga, *et al.* 2014(*28*) generated chromosome-specific graphical models of the α-satellite arrays and reported a bimodal distribution in estimated chrX-specific α-satellite array (DXZ1) sizes (*29*) for a subset of the 1000 Genomes males. Fig. 1b extends this observation to the entire data. The cumulative distributions of estimated array sizes of the three common chrX cenhaps designated in Fig. 1c show substantial differences (Fig. 3a). α-satellite array sizes in cenhap-homozygous females are parallel to males, and imputed cenhap heterozygotes are intermediate, as expected. Similarly, Fig. 3b shows an even more striking example of variation in array size between cenhap homozygotes on chr11, and Fig. 3c demonstrates that heterozygotes of the two most common cenhaps are reliably intermediate in size. While we confirmed that reference bias does not explain the observed cenhaps with large array size on chrX and chr11 (see Methods, Fig. 1b, Fig. 3b and Fig. S4), it is a potential explanation for particular instances of cenhaps with small estimated array sizes, e.g., the relatively low chrX-specific α-satellite content in the highly diverged African cenhap (see Fig 1b,c and Fig. 3a, cenhap 1, highlighted in purple). Importantly, our results demonstrate that cenhaps do robustly tag a substantial component of the genetic variation in array size.

**Figure 3.**
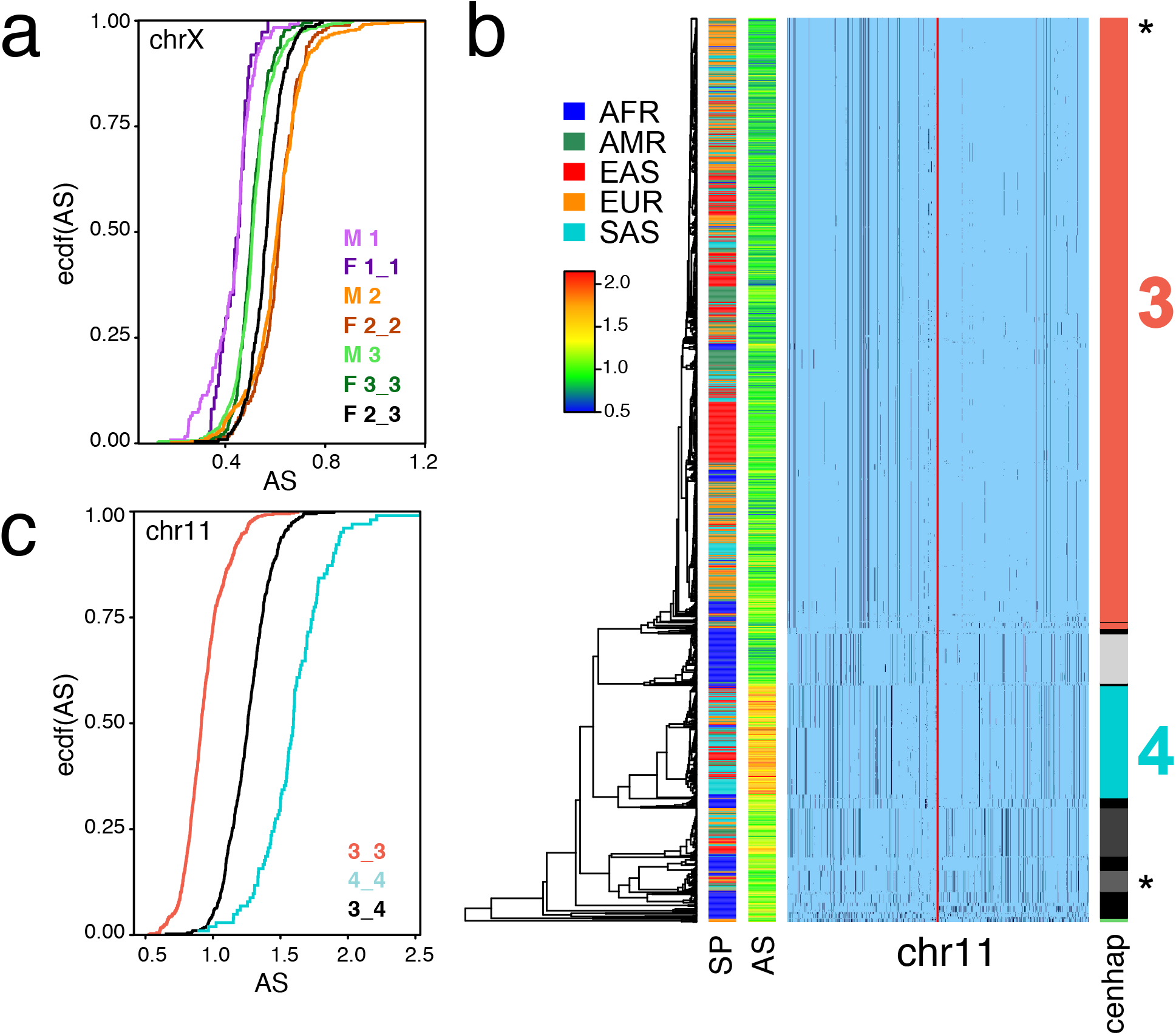
Cenhaps differ in α-satellite array size. **a**. Empirical cumulative density (ecdf) of chrX α-satellite array size for cenhap, homozygotes and heterozygotes. 1_2 and 1_3 heterozygotes were excluded due to insufficient data. Female (F) values were normalized (x 0.5) to facilitate plotting with hemizygote male (M) data. **b**. Haplotypic representation of 1000 SNPs from 1640 imputed chr11 genotypes from 820 cenhap-homozygous individuals. SNPs were filtered for MAC≥35 and passing the *4gt_dco.* Minor alleles shown in black. Assembly gap indicated by red line. Superpopulation (SP) and scaled chr11-specific *α*-satellite array size (AS) at left. Cenhap partitions at right; most common cenhap (‘3”) and cenhap with larger mean array size (“4”) are highlighted. Most probable HuRef cenhap genotypes are indicated by black asterisks at right. c. Empirical cumulative density of array size for chr11 cenhap (from b) homozygotes (3_3 and 4_4) and heterozygotes (3_4).

The potential impact of sequence variation in CPRs and their associated satellites on centromere and heterochromatin functions has been long recognized but difficult to study (*10*). Both binding of the centromere-specific histone, CENPA (*30*) and kinetochore size (*8*) are known to scale with the size of arrays and to fluctuate with sequence variation in satellite DNAs (*7*). Through these interactions with kinetochore function and other roles for heterochromatin in chromosome segregation (*3,31*), α-satellite array variations can affect mitotic stability in human cells (*32*), as well as meiotic drive systems in the mouse (*11*). Meiotic drive has been cited as the likely explanation for the saltatory divergence of satellite sequences and the excess of nonsynonymous divergence of several centromere proteins, some of which interact directly with the DNA (*12*). However, the high levels of haplotypic diversity and deep cenhap lineages we observe (Fig. S2) conflict with the predictions of a naïve turnover model based on strong directional selection yielding sequential fixation of new driven centromeric haplotypes. The inherent frequency-dependence of meiotic drive (*34*), associative overdominance (*33*), a likely tradeoff between meiotic transmission bias and the fidelity of segregation of driven centromeres (*13*), and the expected impact of unlinked suppressors (*34*), are plausible explanations for the surprising levels of cenhap variation.

The identification of human cenhaps raises new questions about the evolution of these unique genomic regions, but also provides the resolution and framework necessary to quantitatively address them. Our results transform large, previously obscure and shunned genomic regions into genetically rich and tractable resources, revealing unexpected diversity and immense archaic centromere introgressions. Most importantly, cenhaps can now be investigated for associations with variation in evolutionarily important chromosome functions, such as meiotic drive (*35*) and recombination(*14*), as well as disease-related functions, such as aneuploidy in the germline(*9*) and in development (*4*), cancer and aging(*3*).

## Acknowledgments

We thank Benjamin Vernot for help accessing archaic DNA sequence data. And we that Graham Coop, Mikkel Schierup and Yuh Chwen Grace Lee for helpful discussions.

## Funding

This work was supported by grants (R01 GM117420 and R01 GM119011) to G.H.K.

## Author contributions

S.A.L and C.H.L. conceived the study and conducted the analyses and wrote the first draft of manuscript. All authors discussed and commented on the manuscript.

## Competing interests

No competing interests declared.

## Data and materials availability

All data needed to evaluate the conclusions in the paper are present in the paper or the supplementary materials.

